# Island-specific evolution of a sex-primed autosome in the planarian *Schmidtea mediterranea*

**DOI:** 10.1101/2021.05.24.445457

**Authors:** Longhua Guo, Joshua Bloom, Daniel Dols Serrate, Eyal Ben David, Olga T. Schubert, Kaiya Kazuma, Katarina Ho, Yubao Wei, Daniel Leighton, James Boocock, Tzitziki Lemus Vergara, Marta Riutort, Alejandro Sánchez Alvarado, Leonid Kruglyak

**Author notes:** To whom correspondence should be addressed. (LG), (LK).

## Abstract

The sexual biotype of the planarian *Schmidtea mediterranea* is a hermaphrodite indigenous to Tunisia and several Mediterranean islands. Here, we isolated individual chromosomes and used sequencing, Hi-C and linkage mapping to assemble a chromosome-scale genome reference. The linkage map revealed an extremely low rate of recombination on chromosome 1. We confirmed suppression of recombination on chromosome 1 by quantifying recombination events in individual sperm and oocytes. We showed that the extensive heterozygous regions, previously designated as J and V haplotypes, comprise essentially all of chromosome 1. Genome sequencing of individuals isolated in the wild indicated that this heterozygosity has evolved specifically in populations from Sardinia and Corsica. We found that chromosome 1 acquired many genes that determine the development of female and male reproductive systems, along with haplotype-specific expression of such sex specific genes. These molecular signatures and restricted recombination in a hermaphrodite led us to propose that chromosome 1 is a sex-primed autosome, providing direct molecular evidence for the traditional model of how sex chromosomes may have evolved from autosomes.

---

Sex chromosomes are thought to have evolved from homologous autosomes that acquired sex-determining genes and lost their ability to recombine (*1–10*). As such, sex chromosome evolution and recombination suppression are closely associated(*1–10*). Yet, direct evidence of such homologous autosomes primed for evolution into sex chromosomes is hard to capture, and little is presently known about the molecular signatures associated with the evolution of recombination suppression.

The freshwater planarian *S. mediterranea* is an important model organism for studies of regeneration (*11, 12*). It exists as asexual and sexual reproductive biotypes. The sexual biotype is mostly distributed in Tunisia, and on the islands of Sardinia, Corsica, and Sicily (*13*). The sexual biotype is a simultaneous hermaphrodite: it develops both male and female reproductive systems in the same adult individual and obligately outcrosses to fertilize other individuals (*14, 15*). We hypothesized that studying chromosome evolution in a simultaneous hermaphrodite might provide insights into the early evolution of a primitive sex chromosome.

*S. mediterranea* has 4 pairs of chromosomes, which are stably diploid. The genome is reported to comprise 774 Mb assembled as 481 non-contiguous series of genomic sequences or ‘scaffolds’ (*16–18*). In a previous study, we found that ~300 Mb of the genome remains heterozygous even after extensive inbreeding of laboratory strains, and that this phenomenon also occurs naturally in wild populations in Sardinia (*14*). The two sets of heterozygous alleles were collectively named the J and V haplotypes. To define the chromosomal locations of these alleles and investigate the reasons underpinning the persistence of heterozygosity in *S. mediterranea*, a detailed assembly of all four chromosomes was needed.

## Chromosome-Scale Genome Assembly

To transform the 481 scaffolds (*16*) into a chromosome-scale genome reference, we carried out chromosome sequencing (ChrSeq) (*19, 20*) of a lab strain, as well as chromatin proximity sequencing by Hi-C (*21, 22*) (**Fig. 1A**). To do so, we dissected individual chromosomes from mitotic cells by using laser capture and amplified and sequenced each chromosome individually (**Fig. 1B**). We examined the sequencing depth of each scaffold across multiple samples of the same chromosome to ensure reproducibility and multiple samples of different chromosomes to ensure specificity (**fig. S1**). Overall, we successfully amplified and confidently assigned to one of the four chromosomes 740 Mb of the 774 Mb genome (**table S1**).

**Figure 1.**
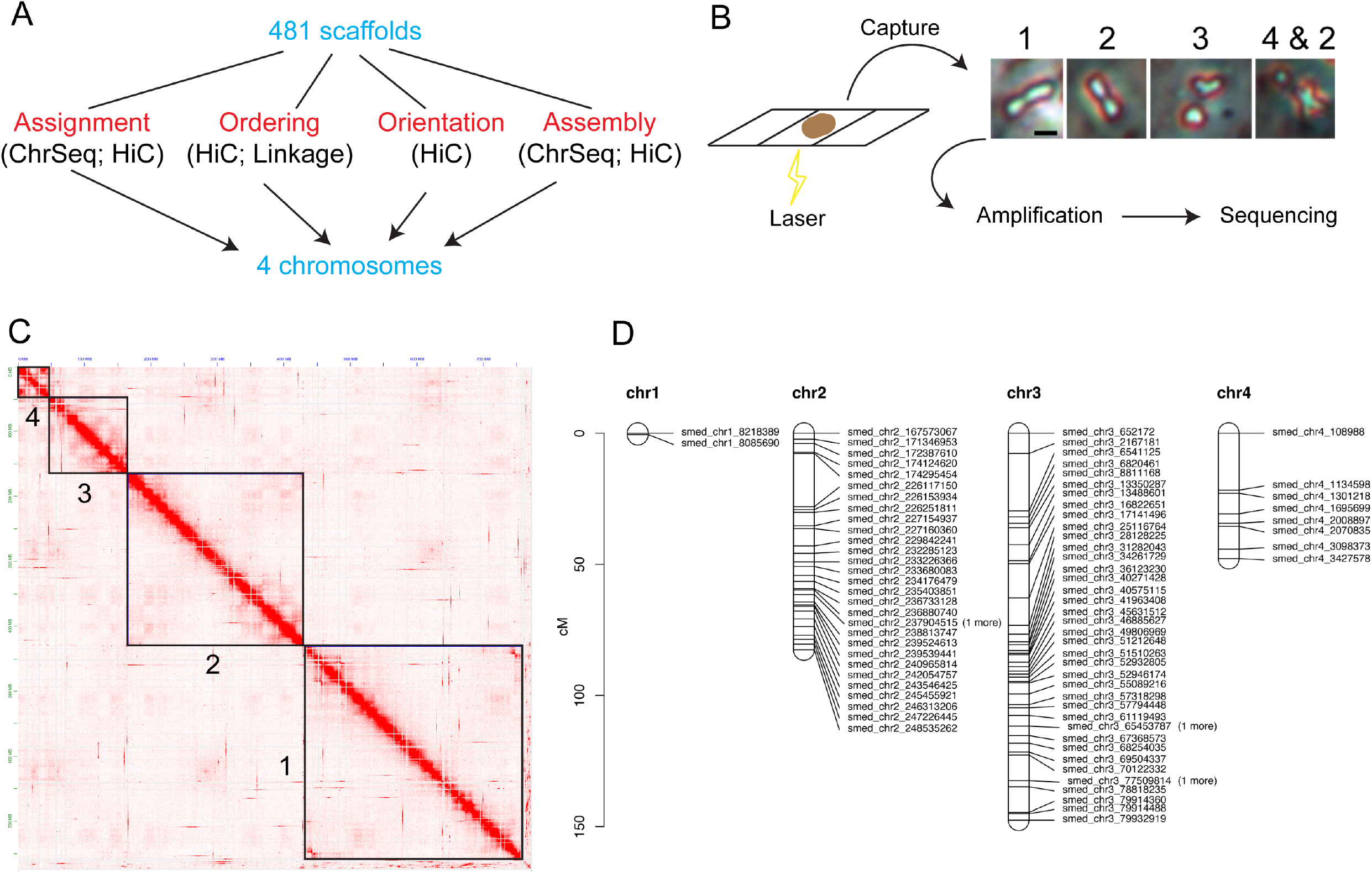
Chromosome-scale genome assembly. (A) Schematic of using Hi-C, ChrSeq, and a linkage map to transform 481 scaffolds (*16*) into a chromosome-scale genome assembly. (B) Chromosomes from mitotic cells were spread onto membrane slides for laser capture and sequencing. Numbers denote the identity of chromosomes present in representative samples. (C) Contact heatmap of chromatin interactions visualized in Juicebox (*23*); black boxes denote the four chromosomes in the final assembly. (D) Linkage map visualized by LinkageMapView (*62*) shows lack of recombination on chromosome 1. Tick marks and labels denote genetic markers.

We used ChrSeq information and chromatin interaction data generated by Hi-C to correct and connect the individual scaffolds within a chromosome into a chromosome-scale genome, hereafter named Smed_chr_ref_v1. Hi-C data analyzed by the scaffolding algorithm SALSA (*21*) resolved 481 scaffolds into 57 super-scaffolds and 104 singletons. ChrSeq revealed that 3 of the 57 super-scaffolds were disjointed inter-chromosomal fragments, consistent with the Hi-C contact heatmap (**Fig. 1C**). We split the disjointed super-scaffolds, merged the scaffolds into chromosomes, and ordered and oriented all scaffolds within the chromosomes by using the visualization software Juicebox (*23*) (**Methods**). Chromosome assignments by ChrSeq alone (**table S1**) and by Hi-C alone were inconsistent for only 3 of 384 (0.8%) scaffolds (**table S2**). We assigned these 3 scaffolds to chromosomes manually based on the Hi-C data. Hi-C also detected 26 inter-chromosomal or intra-chromosomal assembly errors (**table S2**) in the previous assembly (*16*), 5 of which were confirmed by ChrSeq to be inter-chromosomal disjointed scaffolds (**table S2**). The final genome assembly (Smed_chr_ref_v1, **Fig.1C**) has 4 chromosomes with a total size of 764 Mb, 98.4% of that reported previously (*16*). Of the 1.6% of the previous assembly that are not contained in Smed_chr_ref_v1 by Hi-C, around half (52.6%) of the scaffolds could not be assigned to a chromosome or were assigned to 2 chromosomes by ChrSeq, indicating their lower assembly quality (**table S3**); 33.0% and 12.4% of them had chromatin interaction signals with structurally complex regions of chromosome 1 and 2 (**fig. S2**), respectively, suggesting that they might be alternative assemblies or repetitive sequences. In our new assembly, Smed_chr_ref_v1, the two ends of chromosome 4 are capped by >1,000 copies of the telomere repeat TTAGGG, indicating high assembly quality.

To validate the linearity of the chromosomes in Smed_chr_ref_v1, we generated a linkage map. We crossed two divergent laboratory strains of *S. mediterranea*, S2F10b and D5, to produce an F2 population and genotyped individual worms with RADseq (*24*). Eighty markers that are evenly distributed and genotyped in at least 98% of the F2 segregants (91 of 93) were used to establish 4 linkage groups (**Fig. 1D, table S4**). These 4 linkage groups represent the 4 chromosomes. The ordering of the 80 markers in the linkage map is consistent with Smed_chr_ref_v1, independently supporting the quality of our chromosome-scale genome assembly. This highly contiguous and complete genome assembly and linkage map together facilitate further genetic and epigenetic investigations of the functions of the genome in this model planarian.

## Chromosome 1 Recombination Suppression

We reexamined the heterozygous alleles that define the J and V haplotypes in the newly assembled genome. We found that 87.7% of the genetic markers that remained heterozygous were located on chromosome 1, spanning 333 Mb at a density of 30,148 variants per 10 Mb (**Fig. 2A**). The remaining 12.3% of heterozygous markers were located on the other 3 chromosomes at a density of 3,274 markers per 10 Mb; these likely correspond to differences between highly similar copies of repetitive elements rather than to true polymorphisms (**table S5**). All F2 worms (n=93) in this study were heterozygous for chromosome 1 but not for chromosomes 2–4 (**table S6**), consistent with our previous study (*14*). Hence, we conclude that the J/V haplotypes are on chromosome 1.

**Figure 2.**
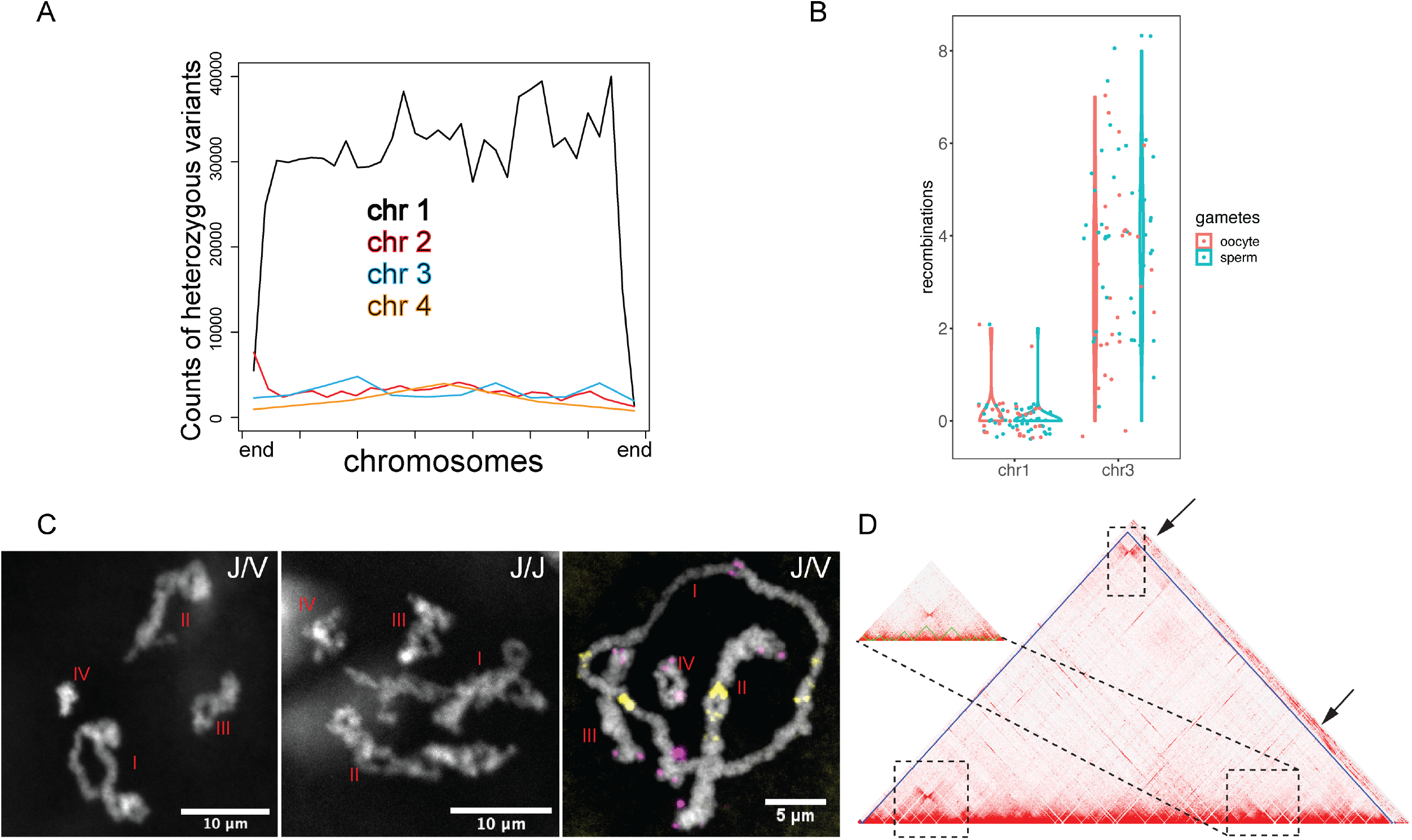
Recombination suppression and inversions on chromosome 1. (A) Distribution of heterozygous variants maintained in the S2 inbreeding pedigree along the four chromosomes; y-axis shows variant counts per 10 Mb. (B) Number of meiotic recombination events on chromosomes 1 and 3 in oocytes (red) and sperm (cyan). Dots represent individual gametes, summarized with violin plots. (C) Chromosome 1 in late prophase I in the oocytes from J/V worms (left and right panels) have fewer crossovers than those in oocytes from J/J worms (middle panel). Chromosomes in the right panel show FISH results with probes for telomeric repeats (TTAGGG, magenta) and centromere-like repeats (yellow). (D) Chromatin contact heatmap for chromosome 1. Dashed-line rectangles indicate locations of potential inversions. Arrows show chromatin contact signals with inversion regions from unassigned scaffolds.

Our linkage map revealed an extremely low rate of recombination on chromosome 1 (only 0.5 cM for the entire chromosome; **Fig. 1D**). This is particularly remarkable because at 333 Mb, chromosome 1 is the largest of the 4 chromosomes, containing more than 40% of the genome.

To examine directly whether chromosome 1 can recombine, we sequenced 45 single sperm and 28 single oocytes from a J/V line, S2 (**Fig. 2B**). Gamete sequencing is preferred because recombination events in hatchlings can be selected for by differential fertilization or embryonic lethality. We identified 3,197 single nucleotide variants (SNVs) on chromosome 1 and 3,312 on chromosome 3, covering 99% of the length of each chromosome (**table S7**). SNVs were distributed at a similar density across 20 Mb windows (coefficient of variation 0.38 and 0.31 for chromosome 1 and 3) (**table S7**). We observed that 98% (44/45) of the sperm and 93% (26/28) of the oocytes had no crossovers on chromosome 1; by contrast, only one sperm and two oocytes had no crossovers on chromosome 3 (**Fig. 2B**). We conclude from this data that recombination on chromosome 1 is strongly suppressed.

Consistent with our conclusion that recombination is suppressed on chromosome 1, we found that during prophase I, when other chromosomes had numerous crossovers, chromosome 1 formed a ring structure in the oocyte in J/V worms but not in J/J worms (**Fig. 2C**). Fluorescence *in situ* hybridization with telomere probes showed that crossovers between the two homologous pairs of chromosome 1 happened only in regions close to the telomeres, leading to the observed ring conformation rather than the side-by-side pairing seen for chromosomes 2, 3 and 4. Furthermore, Hi-C analysis showed that chromosome 1 has three putative inversions, each >20 Mb (**Fig. 2D**); such inversions are an established cause of crossover suppression (*25–27*). The rest of the genome has only one large inversion of ~10 Mb on chromosome 2 (**fig. S2**).

## Island-Specific Evolution of Chromosome 1

To investigate the genetic diversity of *S. mediterranea* in its natural environment and to determine whether chromosome 1 J/V heterozygosity happens in the entire species, we used RADseq (*24*) to sample the genomes of 70 sexual individuals from Sardinia, Corsica, Sicily and Tunisia and 2 asexual individuals from Menorca (*13*) (**Fig. 3A**). To look for genetic relationships between the individuals, we determined clustering of relatedness measured by identity-by-state pairwise distances (*28*) and identified two superclusters: animals from Sicily were closely related to those from Tunisia, and animals from Sardinia were closely related to those from Corsica (**Fig. 3B**). Phylogenetic clustering (**fig. S3**) and a STRUCTURE analysis (**fig. S4**) further supported this population structure.

**Figure 3.**
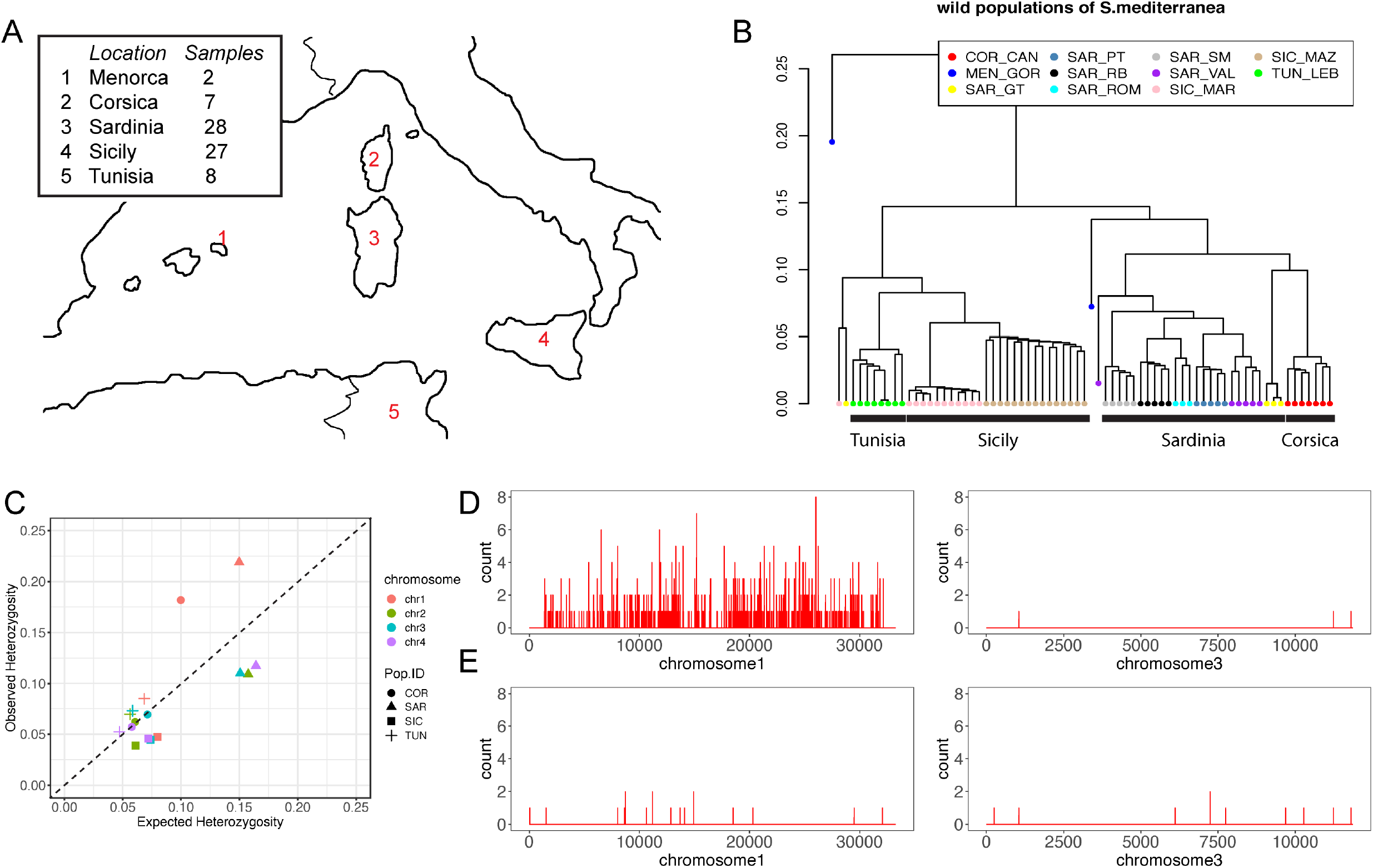
Island-specific evolution of chromosome 1 heterozygosity. (A) Collection sites of 72 wild isolates from Mediterranean islands and Tunisia. (B) Maximum likelihood tree of 2 asexual isolates from Menorca (MEN_GOR) and 70 sexual isolates from Sardinia (SAR), Corsica (COR), Sicily (SIC) and Tunisia (TUN). Dots denote individual worms, with colors corresponding to their collection sites as noted in the legend. (C) Observed vs. expected heterozygosity for each chromosome (denoted by color) in each population (denoted by shape). Dashed line corresponds to equality between observed values and those expected under Hardy-Weinberg equilibrium and is shown as a guide to the eye; the departure of points corresponding to chromosome 1 in COR and SAR from this expectation is notable. (D-E) Number of sites heterozygous in >80% of the population per 10 kb window along chromosome 1 (left) and chromosome 3 (right) in Sardinia (D) and in Sicily (E).

The relatedness of the populations from Sardinia and Corsica suggested that they may share genome characteristics that differ from those of the populations from Sicily and Tunisia. Indeed, animals in the populations from Sardinia and Corsica had greater heterozygosity on chromosome 1 than expected under Hardy-Weinberg equilibrium (**Fig. 3C**), whereas heterozygosity on the other three chromosomes in these populations and on all four chromosomes in the other populations closely followed expectation. By analyzing J/V haplotype markers in the wild populations, we found that the animals from Sardinia/Corsica were heterozygous J/V whereas those from Sicily/Tunisia were homozygous J/J. Moreover, the animals from Sardinia (n=28) contained many sites that were heterozygous in more than 80% of the individuals and that were distributed along the length of chromosome 1 except near the ends, whereas the animals from Sicily (n=27) had very few such heterozygous sites (**Fig. 3, D and E, left**) **(table S8**). Few such heterozygous sites were observed on chromosome 3 in either population (**Fig.3, D and E, right**). These analyses suggest that chromosome 1 specifically evolved and diverged on the islands of Sardinia and Corsica.

## A Sex-Primed Chromosome

To gain insight into the island-specific suppression of recombination on chromosome 1, we examined the genes located on chromosome 1. We found that all currently characterized genes that are essential for development of both male and female reproductive systems were found on chromosome 1, even though differential gene expression analysis between juveniles and adults showed that adult-enriched genes were distributed equally among all 4 chromosomes (**fig. S5**). The critical regulatory genes— *nanos* (*29*), *nhr-1* (*30, 31*), *npy-8* (*32*), *npyr-1* (*33*), and *CPEB-2* (*34*)—were not clustered but distributed along the length of chromosome 1 (**Fig. 4A**). *Nanos, nhr-1* and *CPEB-2* were transcribed from both J and V haplotypes (**fig. S6**). No genetic variants that would allow determination of J- or V-haplotype-specific expression were found in *npy-8* and *npyr-1* transcripts. The presence of these genes on chromosome 1 and their crucial roles in sexual development indicate the importance of chromosome 1 integrity in the maintenance of sexual reproduction. The asexual lineage of *S. mediterranea*, which has a translocation from chromosome 1 to chromosome 3 and is devoid of any reproductive organs(*12*), likely evolved through loss of function of one or several of these genes.

**Figure 4.**
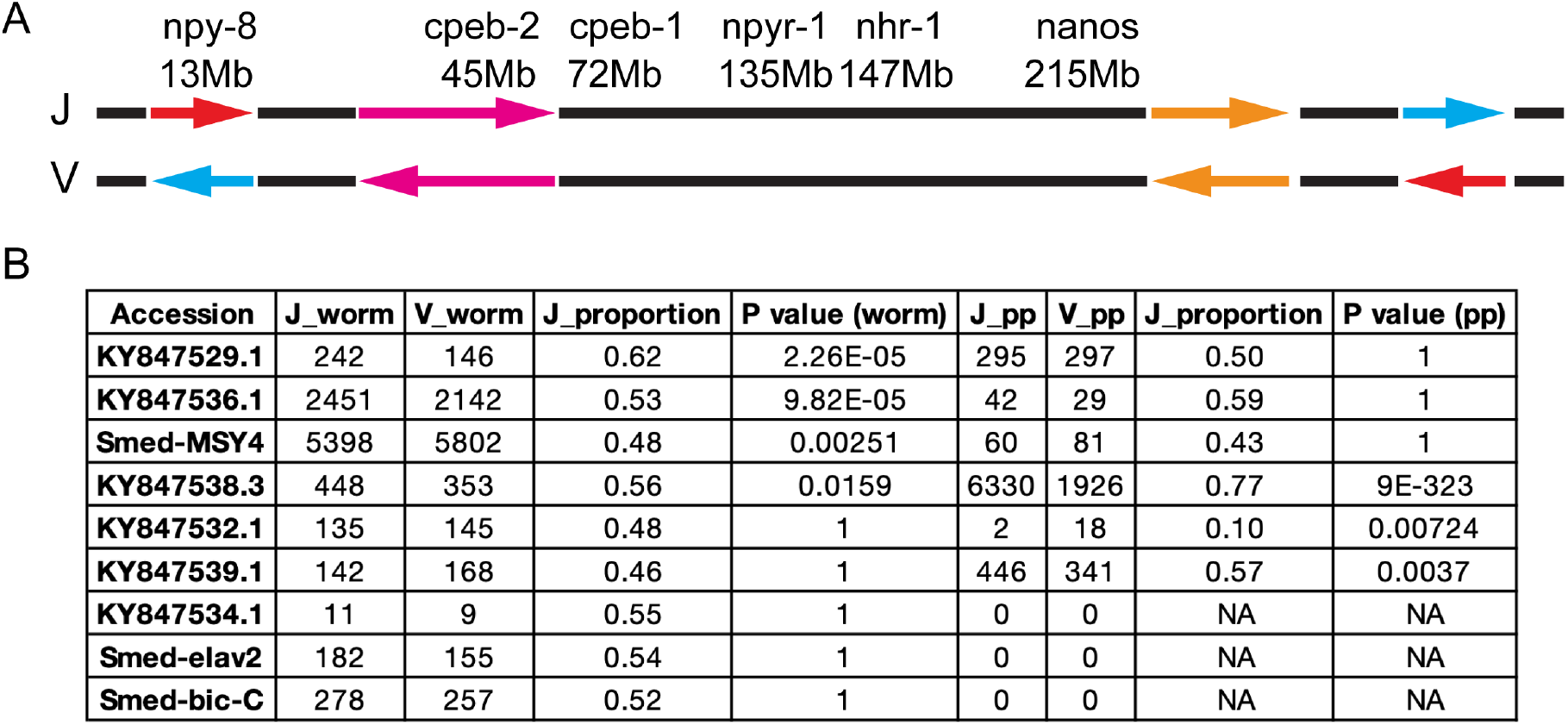
Acquisition and haplotype-specific expression of sex-related genes. (A) A putative model of the structures of the J and V haplotypes of chromosome 1, with locations of critical genes for sexual development (cpeb-1 is female-specific). Arrows denote putative inversions. (B) Comparison of read counts for the J and V alleles of key genes in the transcriptomes of whole sexually mature worms and penis papillae (pp). Bonferroni-corrected P-values from a binomial test of equal expression are shown.

We found that chromosome 1 has acquired many genes with male or female-specific functions. The *CPEB-1* gene on chromosome 1 (**Fig. 4B**) is specifically required for development of the female reproductive system (*34*). The presence of a female-determining gene on a chromosome that does not recombine provides the ideal foundation for the evolution of a sex chromosome: a loss-of-function mutation in the gene on one of the two homologous chromosomes would leave the other chromosome with a unique role in determining the female sex. An extended search of the literature identified 13 additional genes with well characterized male or female-specific functions that are located on chromosome 1 (**table S9**) (*29, 34*). Two of these genes, Bic-C and sped-1, are in synteny with the ZW sex chromosomes of the parasitic flatworm, *Schistosome mansoni* (*35, 36*). Loss of Bic-C expression leads to a “no testes” phenotype, without affecting the ovary (*29*).

To examine whether chromosome 1 has evolved a signature of a sex chromosome— haplotype-specific expression of male- or female-specific genes—we examined the expression of these 14 genes in the transcriptomes of sexually mature adult worms and of their male copulatory organ, the penis papilla (pp) (*37*), in a J/V line. Nine of the 14 genes had at least one heterozygous variant in the coding sequence, which allowed us to determine if these genes showed biased expression from the J or V chromosome. We found that a gelsolin-like gene (KY847538.1) was expressed predominantly from the J chromosome in both whole-worm and pp transcriptomes, with a larger effect in the male copulatory organ. Three genes (KY847529.1, KY847536.1, and Smed-MSY4) had biased expression in the whole worm transcriptome, and two genes (KY847532.1 and KY847539.1) had biased expression in the male copulatory organ (**Fig. 4B, table S9**). Haplotype-specific expression in 6 of 9 female- or male-related genes suggests that sex-specific *cis*-regulatory variation is maintained on chromosome 1.

We have identified a chromosome in the hermaphrodite planarian *S. mediterranea* that does not recombine and has acquired many genes with sex-specific functions, making it primed for evolution into a sex chromosome. The suppression of recombination on this chromosome has evolved specifically on the islands of Sardinia and Corsica; such a phenomenon is absent from the sampled populations of Sicily and Tunisia. Although we cannot know how chromosome 1 will evolve, our findings provide a snapshot that supports the hypothesis that sex chromosomes evolved from homologous autosomes that acquired sex-specific roles and ceased to recombine (*1–8*). The locked J/V heterozygous system may play a facilitating role in this process by maintaining sex-specific alleles in the planarian population prior to the evolution of dioecy (*38*). We propose that the planarian chromosome 1 haplotypes provide a unique opportunity to directly examine the molecular characteristics of a sex-primed autosome.

## Methods

### Chromosome sequencing

Chromosomes were collected from multiple animals of one clonal line, S2, maintained in the lab by amputation and regeneration. Chromosome spreads were prepared on nuclease-free Membrane Slides (Zeiss) according to a previously developed protocol, except that at the last step, the tissues were dissociated into single nuclei and dropped onto the slides without squashing with a coverslip (*39*). Single chromosomes were identified under a 40x lens and collected into caps of single PCR tubes by PALM MicroBeam laser microdissection (Zeiss). Collected chromosomes were spun down with a tabletop spinner in 4*u*l of PBS, and the DNA in the pellets was amplified with REPLI-g Single Cell kit (QIAGEN) for sequencing on the MiSeq or HiSeq 3000 sequencing system (Illumina).

### Chromosome-scale genome assembly

A Hi-C sequencing library was prepared from multiple animals of the S2 strain using the enzyme DpnII, following the instructions of Phase Genomics Proximo Animal kit version 3.0 with few modifications. Sequenced reads were aligned to dd_Smes_g4.fasta (*16*) with bwa mem (version 0.7.17) (*40*). An assembly file was prepared from the SALSA (*21*) output FINAL.fasta with juicebox_scripts (Phase Genomics). The assembly and .hic files were loaded into Juicebox (*23*) for scaffold manipulation (i.e., split, merge, order, orient) and chromosome assembly. The modified assembly file (chromosome-scale) was converted to fasta (Smed_chr_ref_v1) with juicebox_assembly_converter.py.

### Oocyte and sperm sequencing

Sperm were released from sexually mature S2 strain animals into CMFB (1% BSA) buffer. The cell dissociation solution was dropped onto a slide and visualized under a phase contrast microscope to identify single sperm. Oocytes were released from egg capsules. Single sperm or oocytes were transferred into single PCR tubes for amplification with the REPLI-g Single Cell kit (QIAGEN). RADseq libraries and whole genome libraries were prepared and sequenced on the HiSeq 3000 or NovaSeq S2 sequencing system (Illumina). RADseq sequencing data were analyzed as described in the linkage map section. Whole-genome sequencing data were analyzed as described in the recombination section.

### Linkage map

A J/J line, D5, was crossed to a J/V line, S2F8b, to build an F2 population of 93 animals. Genomic DNA was extracted from a fragment of each animal with Easy-DNA gDNA purification kit (K180001, ThermoFisher). Sequencing libraries for RADseq were prepared according to the procedures of Adapterama III(*24*) with few modifications. Genetic variants were identified with Stacks (version 2.41)(*41, 42*). All variants were filtered with VCFtools (version 0.1.14)(*43*) to remove insertions and deletions and to select biallelic single nucleotide variants. Clusters of markers that are located within 200bp were removed as they most likely correspond to repetitive elements. Markers that disobey Mendelian segregation were also removed. Only markers that are homozygous in both parents were used to build the linkage map with R/QTL(*44*).

### Quantifying recombination

Sequencing reads from S2, its oocytes, and sperm were aligned to Smed_chr_ref_v1 with bwa mem (version 0.7.17). Genetic variants were jointly called by using the genome analysis toolkit GATK (v.4.1.4.1) with GenomicsDB and GenotypeGVCFs (*45*). Biallelic heterozygous markers in S2 were further filtered by removing abnormal markers: markers with no segregation of two alleles in the gametes, clusters of markers in close proximity (<200bp), and markers heterozygous in sperm. J and V haplotypes were manually phased with oocytes without recombination and phased by MPR.genotyping (*46*) with all gametes. The MPR.genotyping package was also used to impute or correct missing or erroneous genotypes. The final genotype bins were used to identify and visualize recombination with customized R code. Quantification of recombination was focused on crossovers between long tracks of haplotypes along a chromosome. Putative gene conversion events (i.e., short tracks of haplotype switches encompassing < 1% of chromosome length) were not included.

### Synteny analysis

Protein sequences of genes of interest were obtained from *S. mediterranea*. Such genes were aligned to *S. mansoni* genomes (*36, 47*) with tblastn.

### Gene expression analysis

To examine gene contents on chromosome 1, transcriptome data were downloaded from NCBI Sequence Read Archive (SRA). To examine genes related to sexual development, transcriptomes from sexual adults (*30, 34, 48*), sexual juveniles (*30*), and sexual adults with nhr-1 RNAi (*30*) were used. To examine stem-cell-enriched genes, transcriptomes from sorted X1 cells and CIW4 were used (*49, 50*).

All sequencing data were aligned to dd_Smed_v6 (*17*) with bwa mem (version 0.7.17). Differential gene expression was analyzed with DEseq2 (version 1.26.0) (*51*). Expression was quantified at the transcript level with kallisto (version 0.44.0) (*52*), and imported and summarized to gene-level count matrices by tximport (*53*).

### Haplotype-specific expression

To examine haplotype-specific expression of critical regulators of the reproductive system, mRNA was extracted from six sexually mature animals of a J/V line, S2, and analyzed as three biological replicates, with 2 animals pooled into each replicate. Nine penis papillae (pp) were dissected from nine sexual adult animals of the same line and analyzed as three biological replicates, with pp from 3 animals pooled into each replicate. All mRNA samples were extracted on the same day and processed at the same time for library preparation and sequencing to minimize technical variation. Libraries for RNA-Seq were prepared with Clonetech SMARTer Stranded Total RNA-Seq (Pico) Kit. The workflow consists of converting total RNA to cDNA, and then adding adapters for Illumina sequencing through PCR. The PCR products are purified, and then ribosomal cDNA is depleted. The cDNA fragments are further amplified with primers universal to all libraries. Lastly, the PCR products are purified once more to yield the final cDNA library. Different adaptors were used to multiplex samples in one lane. Sequencing was performed on Illumina Novaseq 6000 with a paired-end-read 150 bp run. Data quality checks were done on Illumina SAV. Demultiplexing was performed with Illumina Bcl2fastq2 v 2.17.

All sequencing data were aligned to dd_Smed_v6 (*17*) with bwa mem (version 0.7.17). To ensure accuracy, haplotype-specific expression of the 14 genes of interest was manually examined in IGV. J or V allele counts were identified for each biallelic variant in the exons. For a particular gene, the allele counts were aggregated from all variants on all exons for J or V haplotypes. The allele counts were then subjected to binomial test and Bonferroni correction (*54*) to determine if the observed allele bias was statistically significant. P-values > 1 after Bonferroni correction were set to 1.

### *De novo* map, parameter optimization and phylogenetic inference

All datasets were run through the *de novo* pipeline as implemented in Stacks version 2.52 (*41, 42*). First, paired-end reads were de-multiplexed and filtered for quality using process_radtags and truncated to a length of 135 bp. Individual reads with Phred-scores below 30 or uncalled bases were discarded (96.9% of reads passed the quality filters). Optimal parameters were identified following Paris et al. (*55*) guidelines by running multiple iterations of the *de novo* pipeline and varying just one parameter with each new iteration on a subset of 13 samples from the same population, [Sic_mar], following recommendations (*56*). We varied the minimum stack depth (-*m*) between 1 and 6 (m1-m6), the number of mismatches allowed between stacks (-*M*) between 1 and 10 (M1-M10), and the number of mismatches allowed to merge catalog loci (-*n*) while keeping all other parameters constant (m3, M2 and n0). We then compared the number of polymorphic assembled loci across samples using the 80% sample representation cutoff (*r80*) and the gain or loss of polymorphic loci with each new iteration. Once -*m* and -*M* were optimized, we assessed -*n* by evaluating the change in number of polymorphic loci for *n* = *M* - 1, *n* = *M* and *n* = *M* + 1. RAD loci were then assembled using the wrapper ‘denovo_map.pl’ in Stacks and the following parameters set: *m* = 3, *M* = 2 and *n* = 3.

Assembled loci that were present in 75% of all individuals were kept from populations, and a minor allele frequency (MAF) filter of 0.04 (--*min-maf*) was used to filter out singleton SNPs that could mask population structure, along with a maximum observed heterozygosity (--*max-obs-het*) filter of 0.99 to remove potentially paralogous loci (*57*). Additionally, to build the phylogenetic tree, we concatenated all RADseq loci after filtering (--*phylip-var-all*). Phylogenetic trees were built by maximum likelihood using RAxML-NG v0.9.0 (*58*), starting from a random seed and applying a GTR+G substitution model and 1000 bootstrap replicates. The sample from Menorca (Sme7-5_men) was used as an outgroup to root the tree.

### Population structure analysis

For this analysis we excluded the *outgroup* sequence and ran populations to retain loci present in all populations (-p 10) and 75% of the individuals present in each population (- r 0.75). Based on the loci that passed our filtering criteria, a random whitelist of 1000 loci was generated and run through populations once again, with the same criteria but retaining the first SNP at each locus (--*write-single-snp*). The output was exported in STRUCTURE format, and STRUCTURE 2.3.4 (*59*) was used to infer population structure with 10,000 chains as burn-in and 100,000 MCMC chains with 20 iterations for *K* = [1-11]. The resulting files were run through STRUCTURE Harvester (*60*), and the optimal *K* was determined (*61*).

## Acknowledgements

We thank Drs. Labib Rouhana, Giancarlo Bruni, and Stefan Zdraljevic for insightful discussions, Zain Kashif and Emily Warda for planarian husbandry, and Xinmin Li and the UCLA Technology Center for Genomics & Bioinformatics (TCGB) for next-generation sequencing. We thank Drs. Ying Zeng, Ekaterina Noskova, and Dobrynin Pasha for discussions on demographic history. We acknowledge Life Science Editors for professional editing services. Confocal laser-scanning microscopy was performed at the Advanced Light Microscopy/Spectroscopy Laboratory and the Leica Microsystems Center of Excellence at the California Nano Systems Institute at UCLA with funding from NIH Shared Instrumentation Grant S10OD025017 and NSF Major Research Instrumentation grant CHE-0722519.

## Data availability

The authors confirm that all data underlying the findings are fully available without restriction. All sequencing data will be available from the NCBI SRA database (accession number PRJNA731187). The chromosome scale genome assembly for sexual *Schmidtea mediterranea* is openly available at: https://planosphere.stowers.org/; http://planmine.mpi-cbg.de/planmine/begin.do

## Figure Legends

**Supplemental Figure 1.**
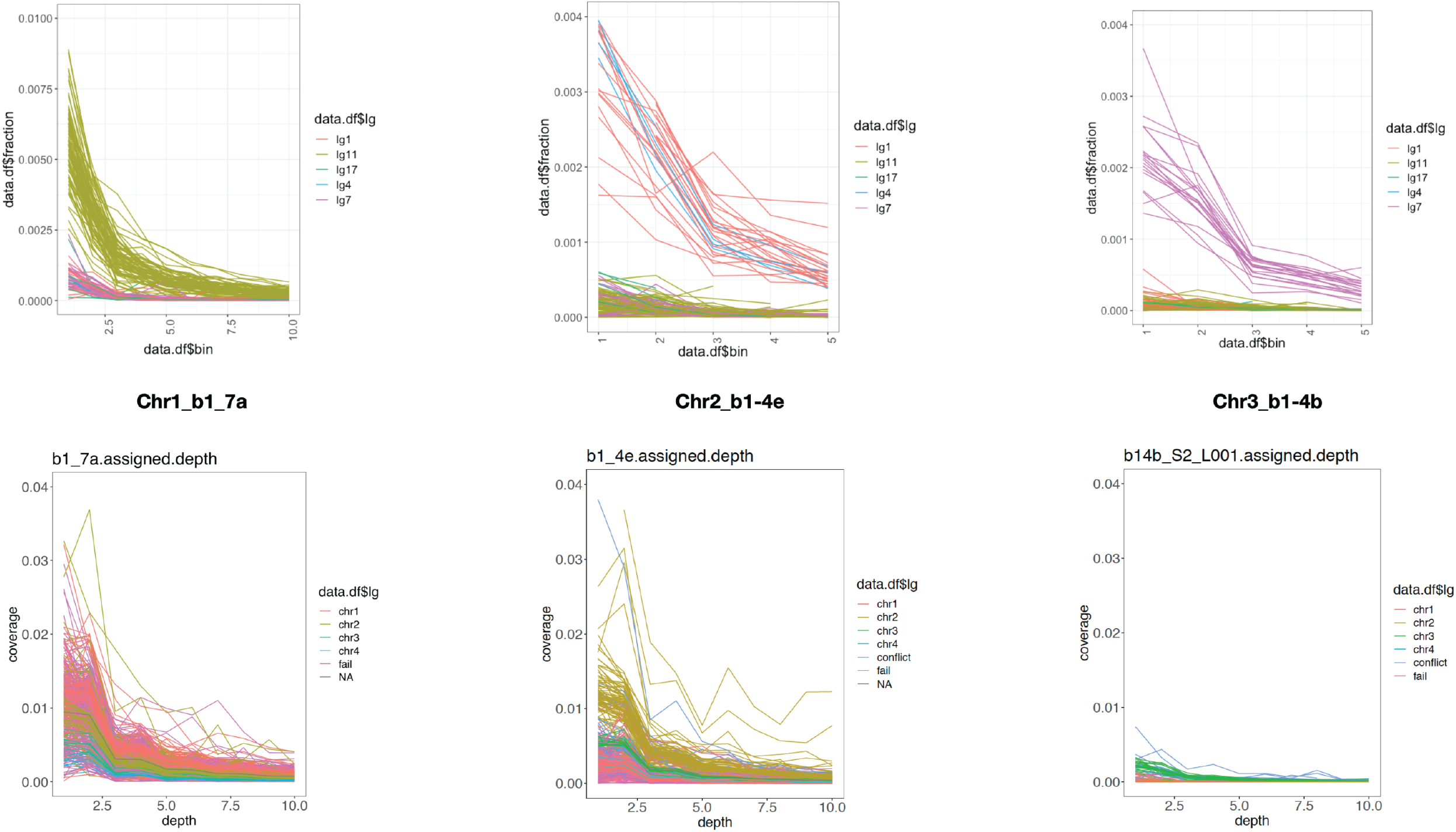
Chromosome assignment by ChrSeq. (A-C) Chromosome sequencing depth of different scaffolds are consistent with linkage group assignment. Linkage groups: lg1, lg11, lg17, lg4, lg7. (D-F) *de novo* assignment of 481 scaffolds to chromosomes by sequencing depth. Chromosomes: chromosome 1 (A, D); chromosome 2 (B, E), chromosome 3 (C, F).

**Supplemental Figure 2.**
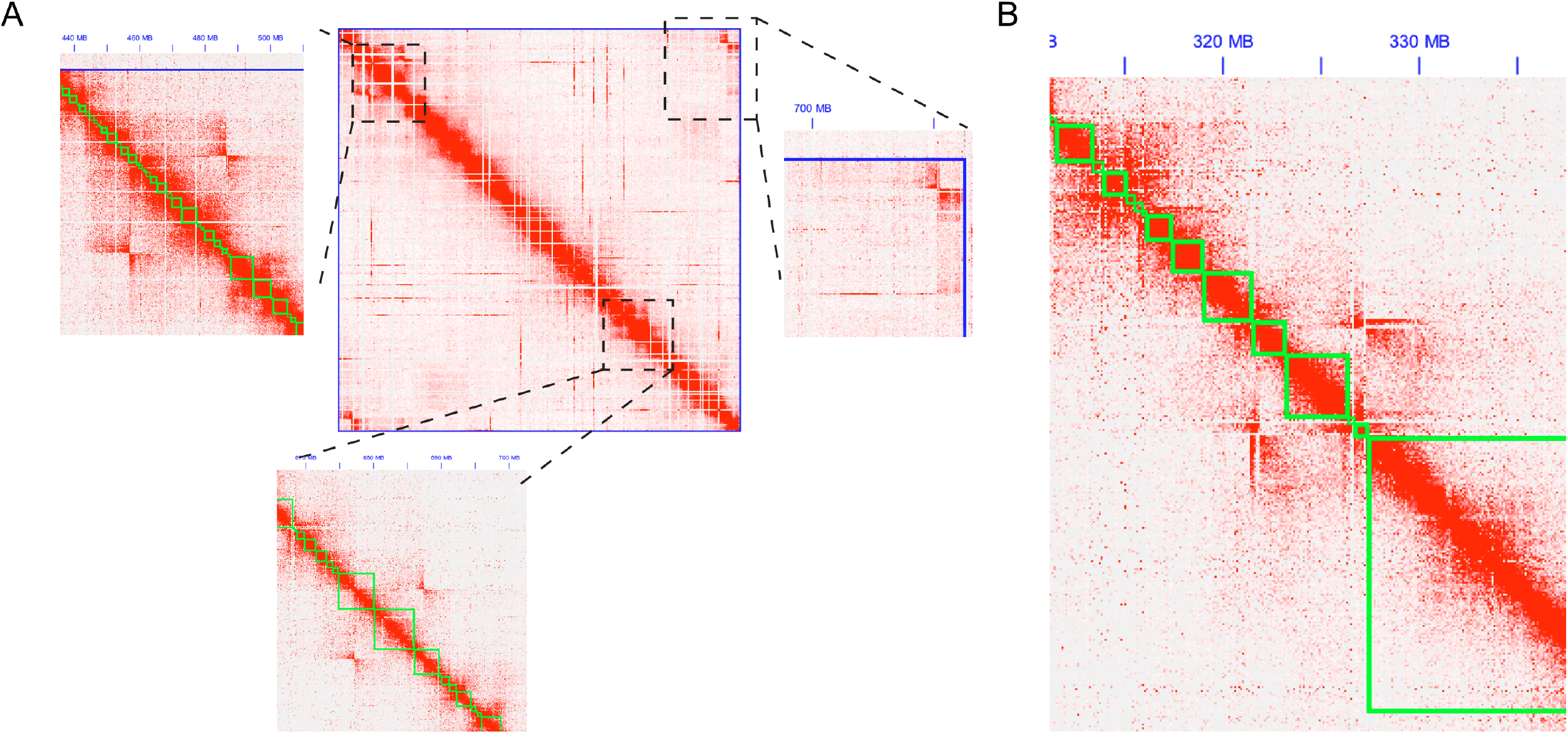
Putative inversions in the planarian genome detected by chromatin contact heatmap. (A) chromosome 1 chromatin contact heatmap showing 3 potential inversions (dashed-line rectangles) visualized in Juicebox. (B) A complex structural variant on chromosome 2. Green outline: dd_Smes_g4 scaffolds. Blue outline: chromosome boundaries.

**Supplemental Figure 3.**
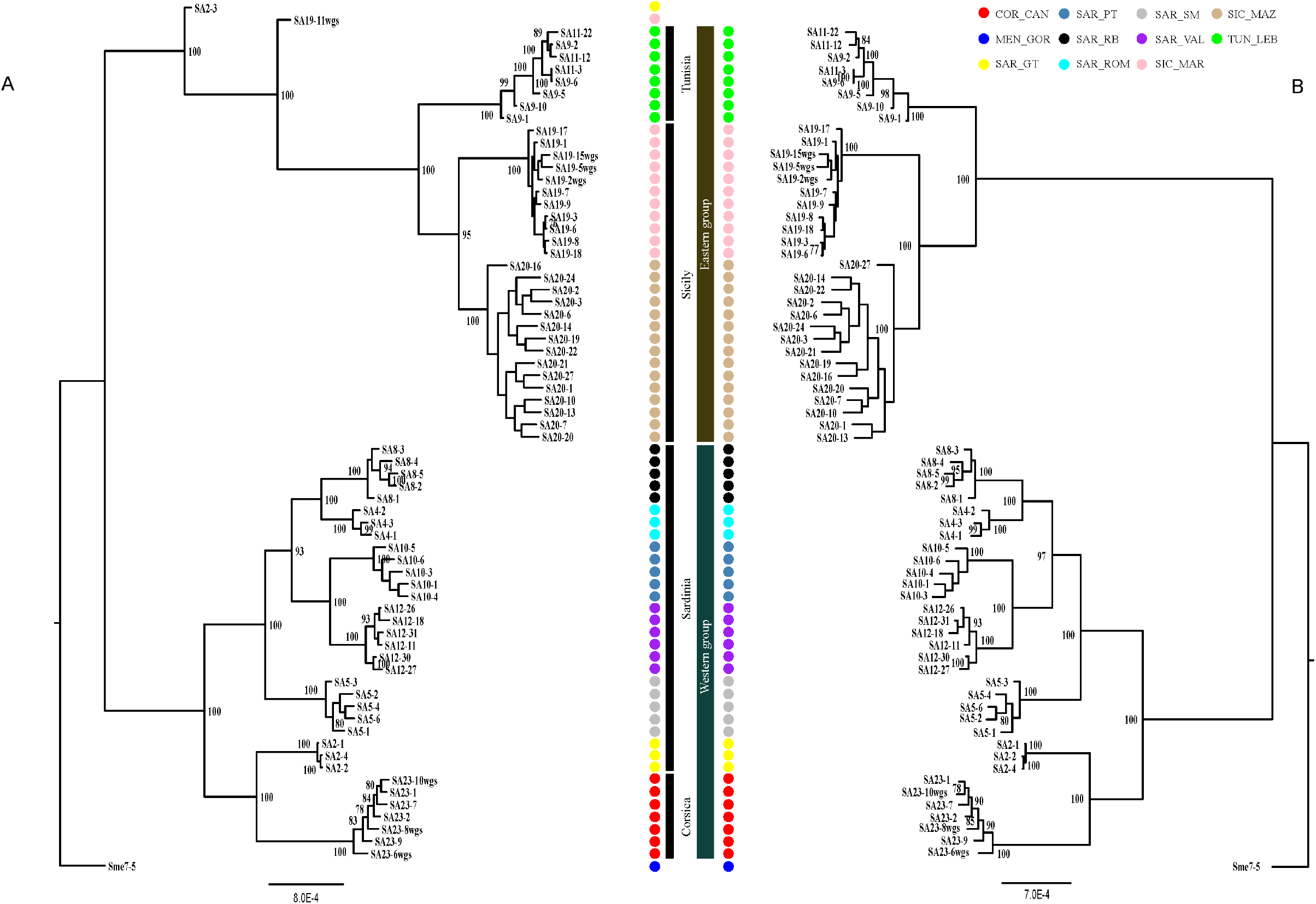
Phylogenetic tree of wild isolates of *Schmidtea mediterranea*. The tree is rooted with one asexual animal from Menorca (Sme7-5). The tree on the left (A) includes two animals from SAR and SIC with higher genome heterozygosity than the cohorts from the same collection sites. Phylogenetic relations agree with the previous designation of eastern and western populations (*13*). SAR: Sardinia; CoR: Corsica; SIC: Sicily; TUN: Tunisia.

**Supplemental Figure 4.**
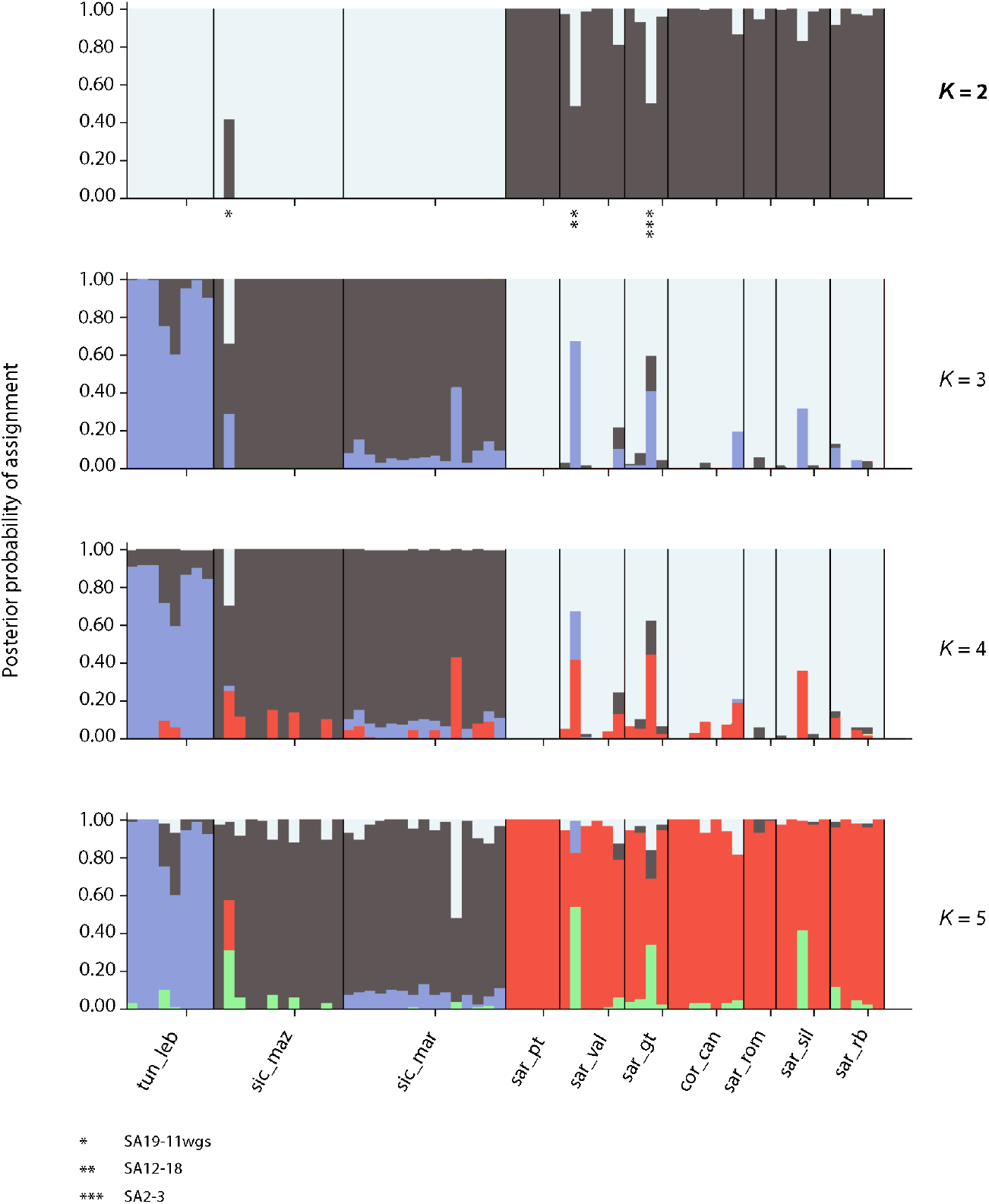
STRUCTURE analysis of wild isolates. Evanno analysis supports a K=2 split, dividing geographically the samples in an Eastern and a Western group. With K=3, a Tunisian signal was found to be present in all other 3 groups (light blue).

**Supplemental Figure 5.**
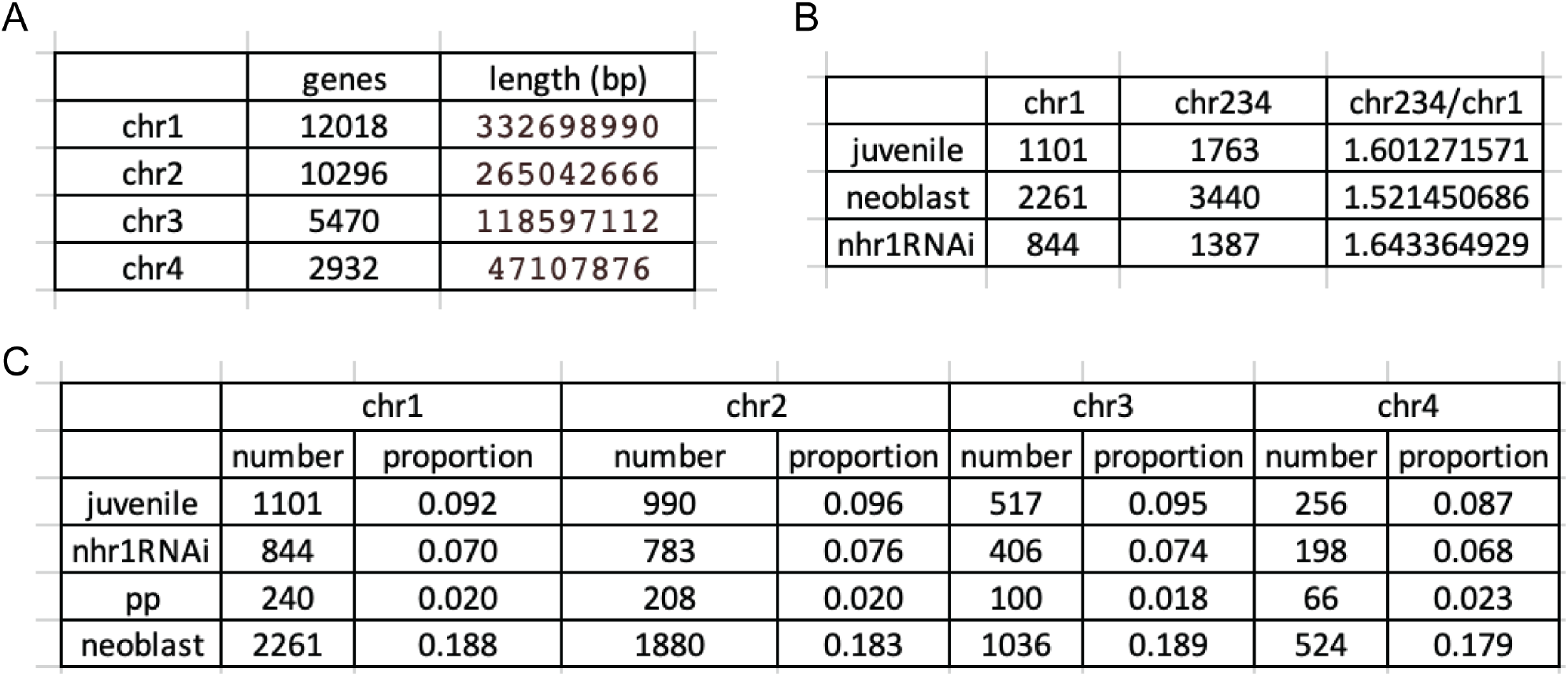
Genes enriched in the sexual reproductive system were distributed equally on all 4 chromosomes. (A) Number of genes and total length of each chromosome. Genes were based on dd_Smed_v6 annotation. (B) Differentially expressed genes and their localizations on chromosome 1 and the rest of the chromosomes. chr234/chr1: the ratio of genes on chr2, 3, and 4 to chr1. (C) Differentially expressed genes and their localizations on chromosome 1, 2, 3, and 4. Proportion: number of differentially expressed genes to the total number of genes per chromosome. (B-C) juvenile: comparing public transcriptome of juvenile animals to adult animals (*30, 34, 48*); nhr1 RNAi: comparing public transcriptome of nhr-1 RNAi animals to adult animals (*30*); pp: comparing transcriptome of penis papillae to whole worm, generated in current study; neoblast: comparing public transcriptome of sorted X1 cells to whole worm in CIW4 animals (*49, 50*).

**Supplemental Figure 6.**
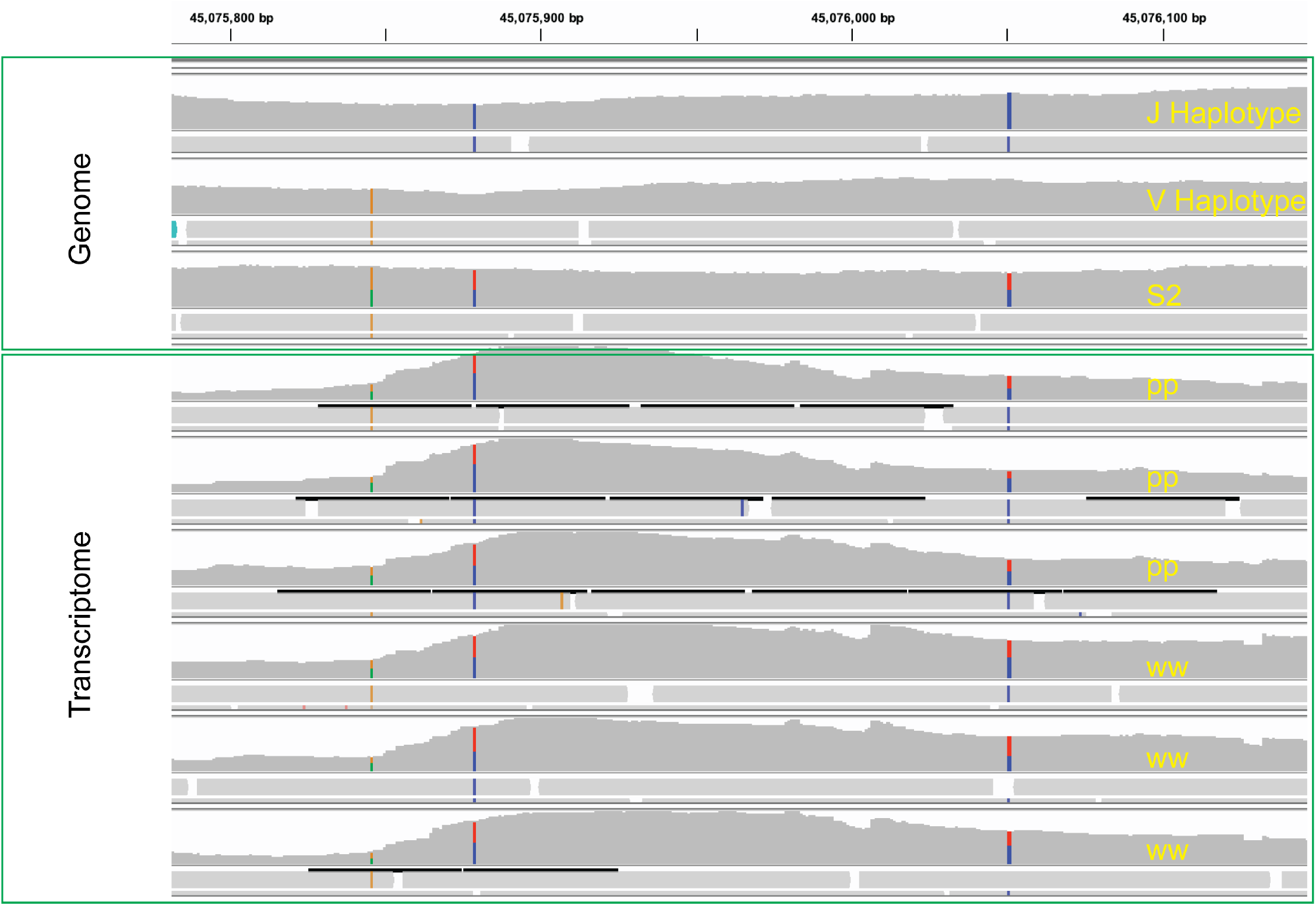
*Nanos, nhr-1* and *CPEB-2* transcribed from both J- and V-haplotypes. The sequencing reads from genomes (J haplotype, V haplotype, S2) and transcriptomes (pp, ww) were aligned to Smed_chr_ref_v1. Shown above are reads for the gene CPEB-2. Three variants with J and V alleles are marked by colored bars. In the genome of the adult worm S2, both J and V alleles are present at equal ratios. In the genome of oocytes (J and V haplotype), the variants have only one allele. In the transcriptomes of penis papillae (pp) and whole worm (ww), both J and V alleles are present at equal ratios.

**Table S1 Assigning 481 scaffolds to chromosomes with 17 sequenced chromosomes**

Chromosome sample information (i.e., ploidy, identity and potential contamination), sequencing coverage and the chromosome assignment for each of the 481 scaffolds from the assembly dd_Smes_g4. In chromosome assignment, 0 (scaffolds that could not be assigned to chromosomes by chrseq); 5 (scaffolds that were assigned to two chromosomes by chrseq); 1-4 (chromosomes).

**Table S2 Linking 481 dd_Smes_g4 scaffolds into chromosomes with Hi-C chromatin contact sequencing**

dd_Smes_g4 scaffolds that were assigned or unassigned to Smed_chr_ref_v1 (chrAssembly), their identity in SALSA assembly of raw Hi-C sequencing data (salsa_scaffolds), and their assignment to chromosomes by chromosome sequencing (chrseq). In chromosome assignment, 0 (scaffolds that could not be assigned to chromosomes by chrseq); 5 (scaffolds that were assigned to two chromosomes by chrseq); 1-4 (chromosomes).

**Table S3 Summary of the final chromosome scale genome assembly (Smed_chr_ref_v1)**

Hi-C detected 26 assembly errors in dd_Smes_g4, 5 of which were confirmed by chromosome sequencing to be inter-chromosomal mis-joining. A total of 97 dd_Smes_g4 scaffolds were not assigned to Smed_chr_ref_v1.

**Table S4 Genetic markers and distances in the linkage map established from a F2 mapping population**

Linkage groups L.3, L.6 and L.8 only contained one genetic marker, which were not included in the table. Chromosomes 2 and 4 were split into two linkage groups each, likely due to the small size of the F2 mapping population.

**Table S5 Distribution of Genetic variants that are maintained heterozygous in the inbreeding pedigree**

Chromosomes were divided into 10Mb windows. Heterozygous variants were identified from the inbreeding pedigree from S2 to S2F9b.

**Table S6 Chromosome 1 was heterozygous in all samples of a F2 population**

The genetic markers were heterozygous in the J/V parent (parent_S2F10B_2A and parent_S2F10B_2B) and homozygous in the J/J parent (parent_D5-1). parent_S2F10B_2A and parent_S2F10B_2B are clones. Genotyping data from RADseq of 291 F2 samples were listed. The 291 F2 samples correspond to 93 unique segregants. 0/0: homozygous reference allele; 1/1: homozygous alternative allele; 0/1: heterozygous; ./.: missing data.

**Table S7 Recombination in the gametes**

Distribution of heterozygous variants identified in the J/V line S2 and used in the crossover assessment along chromosome 1 and 3, in 20Mb windows. Number of crossovers identified per gamete on chromosome 1 and 3.

**Table S8 Chromosome 1 genotypes of wild isolates in Sardinia and Sicily**

Genotyping data for 6 different collection sites in Sardinia and 2 different collection sites in Sicily were aggregated. Row 1 are names of individual animals. 0/0: homozygous reference allele; 1/1: homozygous alternative allele; 0/1: heterozygous; ./.: missing data.

**Table S9 Allele specific expression of “male” and “female” genes**

Genes with well characterized sex-related functions were identified from published work. Their locations on Smed_chr_ref_v1 and *Schistosoma mansoni* chromosomes were determined. The expressed J or V alleles for each of the 9 ‘male’ or ‘female’ genes were quantified in the transcriptomes of whole worms and isolated male copulatory organ, penis papillae.

